# Effects of depression on prefrontal-striatal goal-directed and habitual control

**DOI:** 10.1101/381152

**Authors:** Suyeon Heo, Sang Wan Lee

**Affiliations:** Department of Bio and Brain Engineering, Korea Advanced Institute of Science Technology (KAIST), Daejeon 34141, Republic of Korea; Brain and Cognitive Engineering Program, Korea Advanced Institute of Science Technology (KAIST), Daejeon 34141, Republic of Korea; KAIST Institute for Health Science Technology, Korea Advanced Institute of Science Technology (KAIST), Daejeon 34141, Republic of Korea; KAIST Institute for Artificial Intelligence, Korea Advanced Institute of Science Technology (KAIST), Daejeon 34141, Republic of Korea

## Abstract

Depression is characterized by deficits in the reinforcement learning (RL) process. Although many computational and neural studies have extended our knowledge of the impact of depression on RL, most focus on habitual control (model-free RL), yielding a relatively poor understanding of goal-directed control (model-based RL) and arbitration control to find a balance between the two. We investigate the effects of depression on goal-directed and habitual control in the prefrontal–striatal circuitry. We find that depression is associated with attenuated state and reward prediction error representation in the insula and caudate, a disruption of arbitration control in the predominantly inferior lateral prefrontal cortex and frontopolar cortex, and suboptimal value–action conversion. These findings fully characterize how depression influences different levels of RL, challenging previous conflicting views that depression simply influences either habitual or goal-directed control. Our study creates possibilities for various clinical applications, such as early diagnosis and behavioral therapy design.

## INTRODUCTION

Major depressive disorder (MDD) has received considerable attention, as the lifetime prevalence of the disorder is higher than 10% worldwide^1^. MDD is characterized by deficits in decision-making^2,3^ and its underlying reward learning processes^4^. Recently, with the development of computational models, several studies have explored how depression influences the reward learning system (for a review, see (Chen et al., 2015)).

Reinforcement learning (RL), the process of learning to develop a behavioral policy to maximize reward^5^, has been known to be guided by the two distinct RL strategies: model-based (MB) RL and model-free (MF) RL, each of which guides goal-directed and habitual RL, respectively^6^. Model-based RL guides context-sensitive and goal-directed behaviors through a sophisticated process in which the learning agent makes decisions by simulating an internal environmental model, whereas model-free RL is associated with habitual responses to reward-predicting stimuli based on learned associations between stimuli and rewards^6,7^. Mounting evidence suggests that depression is characterized by impairments in either model-based or model-free RL. For example, behaviors in depressive individuals can be accounted for by impaired model-based RL^8–10^ or a transition from model-based to model-free RL^11^. However, most studies have only explored the effect of depression on model-free RL. For instance, depressive people exhibit an impaired ability to learn stimulus–reward associations accompanying inaccurate representations of reward prediction error^12–17^ or abnormal learning rate control^11,18^.

Impairment in RL is associated with not only the onset of depressive symptoms, but also the development of depression. For instance, stress, one of the major risk factors for depression^19,20^, can induce deficits in RL. Previous findings have shown that people exhibit a reduced ability to engage in model-based RL under conditions of chronic^21,22^ and acute^23,24^ stress. These findings suggest a gradual impairment of RL from the very early stages of depression.

Although these studies have contributed to our understanding of depression in the context of RL, it is still unclear whether depression is best characterized by model-free RL, model-based RL, or an interaction between the two, or how depression influences the neural circuits guiding goal-directed and habitual behavioral control. Moreover, little is known about how these cases extend to early or mild depression.

Here, we aim to provide a computational and neural account of how depression affects goal-directed and habitual control in the prefrontal–striatal circuitry. First, to investigate the effects of depression on model-based and model-free RL, we ran a model comparison analysis to identify a version of arbitration control intended to account for various behavioral traits of depression. In particular, our computational models consider sub-optimality of RL, allowing us to explain choice behavior patterns across a wide spectrum of depression. We combine this with model-based functional magnetic resonance imaging (fMRI) to identify the parametric effects of depression on neural systems associated with model-based and model-free RL. In the subsequent analysis, we attempt to fully characterize how depression disrupts the arbitration between model-based and model-free RL by combining the results from the computational modeling and model-based fMRI analyses and the multi-voxel pattern analysis (MVPA).

## RESULTS

### Effects of Depression on Behavior Performance of Goal-directed and Habitual Learning

Sixty-three participants conducted a sequential two-stage Markov decision task^25^. Of the subjects, 28 were scanned with an fMRI while performing the task. The task manipulated both prediction uncertainty and task goals to dissociate goal-directed and habitual behavior control (Figure 1; for more detail, see Methods). The task consisted of four types of blocks (high/low state–action–state transition uncertainty x specific/flexible goal condition). Before each experiment, participants completed the Center for Epidemiologic Studies Depression (CES-D) questionnaire^26^ (For the distribution of participant’s depression severity, see Supplementary Figure S2).

**Figure 1:**
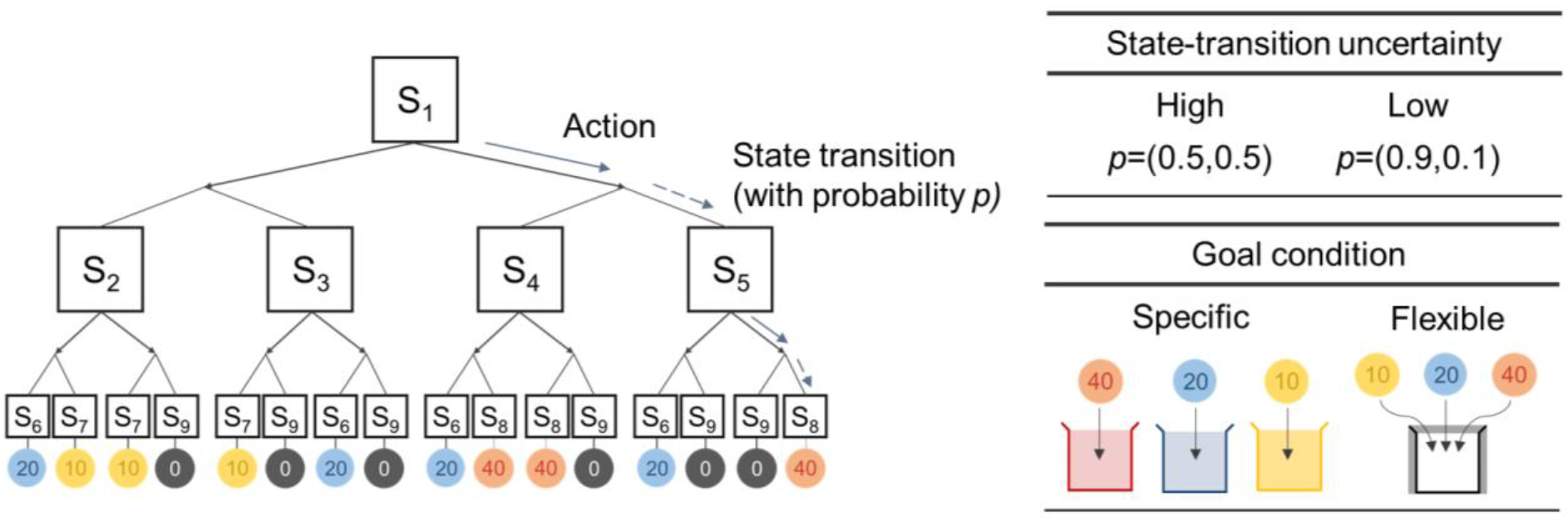
Markov decision task structure. We use the two-stage Markov decision task proposed by Lee et al. (2014). In each stage, participants make a binary choice (left or right). After the first choice in the initial state (S_1_), they were moved forward to one of four states in the second stage (S_2_ - S_5_) with certain state-action-state transition probability *p*. The transition probability (0.5, 0.5) and (0.9, 0.1) corresponds to a high-uncertainty and a low-uncertainty environment, respectively. The task consists of the two goal conditions: a specific-goal and a flexible-goal condition. In the specific goal condition, subjects can collect coins (redeemable for monetary reward) only if the coin color matches with the color of the token box (red, blue, yellow). In the flexible goal condition indicated by the white token box, all types of coins are redeemable.

Overall task performance is negatively correlated with the self-reported depression score. The accumulated reward decreases significantly as individual depression score (CES-D) increases (correlation coefficient estimate=-0.584 [*p*=4.94e-07]; Figure 2a). The proportion of optimal choices, the measure that quantifies the extent to which a subject’s choice reflects an optimal policy, is also inversely proportional to the CES-D score (correlation coefficient estimate=-0.567 [*p*=9.32e-05]; Figure 2b). Finally, choice consistency, the proportion of making the same choice as in previous trials, decreases as the CES-D score increases (correlation coefficient estimate=-0.472 [*p*=1.24e-06]; Figure 2c). These results demonstrate that depression has a damaging effect on both goal-directed and habitual control performance, leading to suboptimal choices.

**Figure 2:**
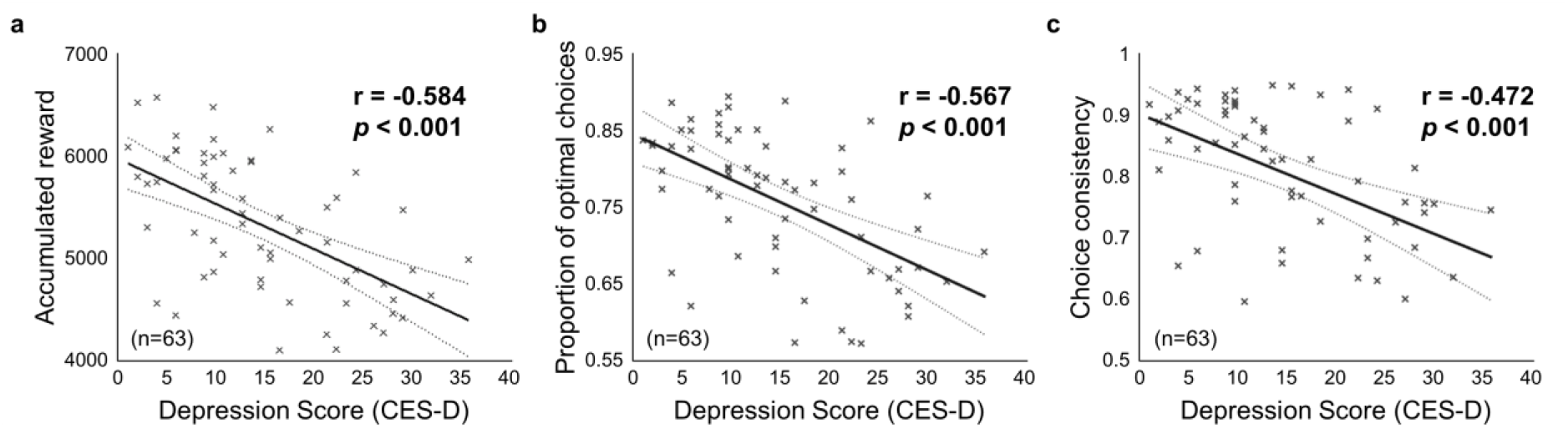
Behavioral results. (a) Relationship between the individual depression score (CES-D) and accumulated reward (n=63). The task performance decreases as the depression score increases. (b) Relationship between depression score and the proportion of optimal choices (n=63). The proportion of optimal choices is inversely proportional to the depression score. (c) Relationship between the depression score and choice consistency in the first state (S_1_) (n=63). The choice consistency index is negatively correlated with the depression score.

### Computational Model of Arbitration Control Allowing for Suboptimal Decision-Making

We adopted the previous dynamic arbitration control hypothesis that respective prediction uncertainty—specifically, the amount of uncertainty in the state and reward prediction error of model-based and model-free RL—mediates the trial-by-trial value integration of the model-based and model-free systems^25^. To fully explore the effects of depression on arbitration control, however, a model should be flexible enough to account for any individual variability arising from suboptimal learning and decision-making.

To consider this, we redesigned the arbitration control scheme to allow for sub-optimality of RL in both learning values and converting learned values into choice behavior. We included the former by introducing separate learning rates for model-based and model-free RL and the latter by defining an exploitation sensitivity parameter as a function of model preference for either model-based or model-free RL (for more detail, see Methods). This model setting reflects the hypothesis that prediction uncertainty mediates not only value integration, but also value-action conversion (Figure 3a).

**Figure 3:**
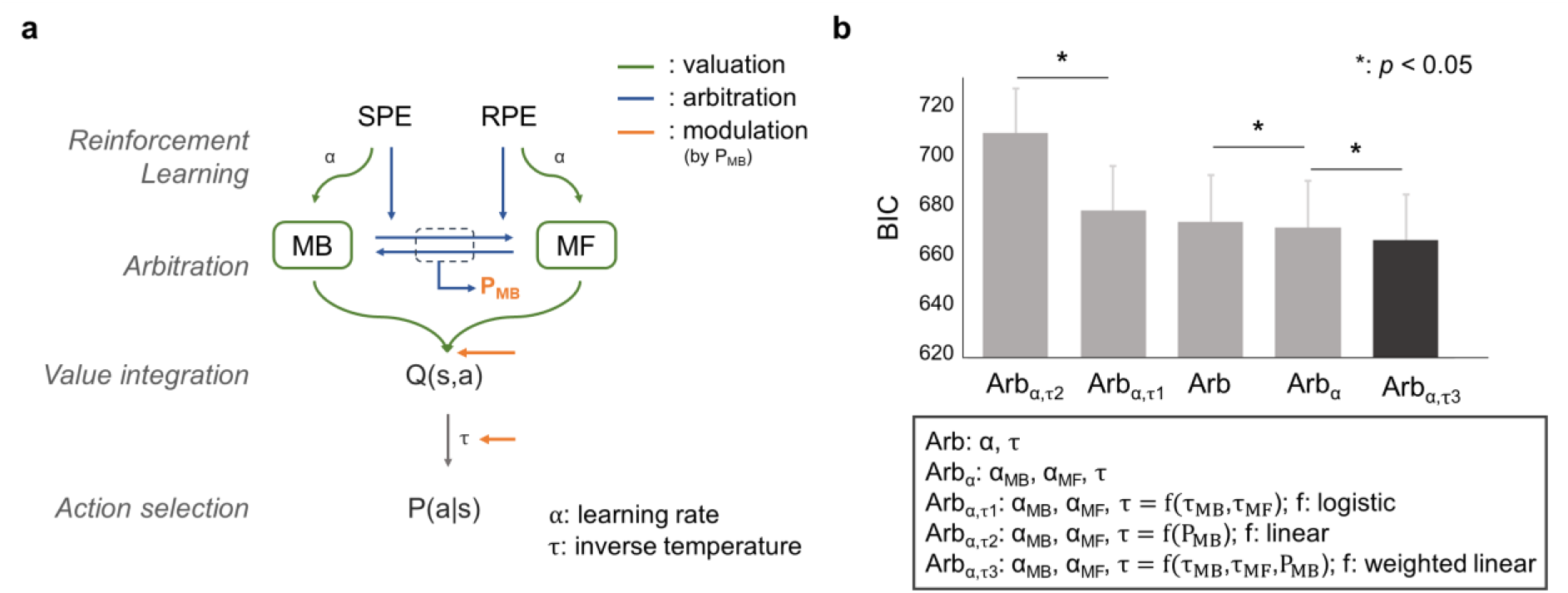
Computational model of dynamic arbitration control. (a) Computational model to investigate dynamic control mechanisms to arbitrate between model-based (MB) and model-free (MF) RL. Two separate RL systems update an action value and reliability of its prediction by using prediction errors (SPE and RPE, respectively; green). The reliability values of the two systems were then used to compute the model choice probability (P_MB_) (blue). The model choice probability guides both the value integration and value-action conversion process; both effects are indicated by the orange arrow. (b) Model comparison analysis. We used Bayesian Information Criteria (BIC) for comparing the goodness of fit while penalizing for the model complexity. Arb refers to the computational model proposed in (Lee et al., 2014). Additional versions of arbitration control consider separate learning rates and dynamic exploitation. Arb_α_ assigns separate learning rates to two different systems (i.e. use α_MB_, α_MF_ instead of α). Arb_α,τ_ is the same as Arb_α_, except for the assumption that the degree of exploitation is a function of the model choice probability. The best version of model uses the degree of exploitation (τ) as a weighted sum of the model-based and the model-free exploitation parameter (τ_MB_ and τ_MF_, respectively) with the model choice probability P_MB_ (i.e. τ = P_MB_*τ_MB_ + (1-P_MB_) * τ_MF_). The error bar stands for the standard error mean.

We compared prediction performance for the five different versions of arbitration control, including the original arbitration model and four other versions implementing our hypothesis in different ways. We used the Bayesian information criteria (BIC) as a performance measure to preclude overfitting. We found that the version implementing our hypothesis, in which the degree of exploitation is determined by the weighted sum of the model-based and model-free exploitation parameters with the model choice probability, explains the subjects’ behavior (*t*(62)=2.46, [*p*=0.017]; paired t-test comparison with the second-best model, Arbα), significantly better than the original model^25^ (Figure 3b; for a model comparison, see Supplementary Table S1; for estimated model parameters, see Supplementary Table S2). Our model comparison result not only corroborates the previous finding that prediction uncertainty mediates the arbitration between model-based and model-free RL, but also demonstrates the effect of prediction uncertainty on both value integration and value–action conversion.

### Goal-directed and Habitual Control in the Prefrontal-Striatal Circuitry

To further examine whether our model explains the neural activity patterns of brain areas previously implicated in model-based and model-free RL, we ran a model-based fMRI analysis in which each of the key signals of our computational model were regressed against the fMRI data.

First, we replicated previous findings concerning the neural correlates of prediction error for the model-based and model-free systems. The state prediction error (SPE) was found bilaterally in the insula and the dorsolateral prefrontal cortex (dlPFC) (all p<0.05 in cluster-level corrected). The reward prediction error (RPE) was correlated with neural activity in both sides of the ventral striatum (p<0.05 in family-wise error [FWE] corrected). These results are fully consistent with previous findings^25,27,28^ (Figure 4, Supplementary Table S3).

**Figure 4:**
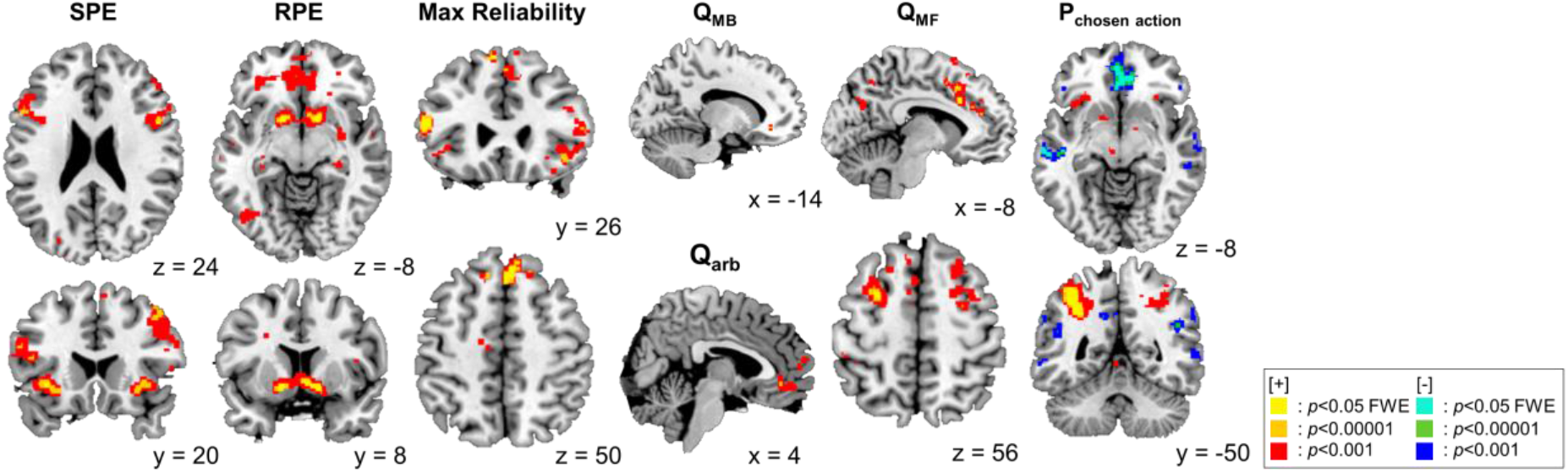
Neural correlates of dynamic arbitration control. Prediction error, value, reliability, action choice probability signals from the proposed model are shown as colored blobs. SPE and RPE refers to state prediction error and reward prediction error, respectively. Max reliability refers to the reliability of whichever system had the highest reliability index on each trial (=max(Rel_MB_, Rel_MF_)). Q_MB_ and Q_MF_ indicate the chosen value from the model-based and model-free system. Q_arb_ refers to the difference between integrated chosen action value and unchosen action value. P_chosen action_ refers to the probability assigned to the chosen action. See more detailed information in Supplementary Table S3-S5.

We also successfully replicated previous findings supporting the neural hypothesis of arbitration control. We found the max reliability signal, the key signal used to mediate arbitration between model-based and model-free RL, in the bilateral inferior lateral PFC (ilPFC, left: p<0.05 in cluster-level corrected; right: p<0.05 in small-volume corrected [SVC] at [8,6,-2]) and the frontopolar cortex (FPC, p<0.05 in cluster-level corrected), fully consistent with previous results^25,29^ (Figure 4, Supplementary Table S3).

Next, we tested the brain areas implicated in value computation. The chosen value of the model-based system (Q_MB_) was found to be encoded in the precentral gyrus (p<0.05 in cluster-level corrected) and the orbital and medial PFC (p<0.05 in SVC at [-14,36,-8])^28,30^. The chosen value of the MF system (Q_MF_) was found in the dorsal ACC, supplementary motor area, premotor cortex, dorsolateral PFC (p<0.05 in cluster-level corrected), and dorsomedial PFC (p<0.05 in SVC at [9,35,40])^31,32^. Notably, this model-free value signal was also found in the posterior putamen, the brain area known to be involved in valuation for habitual learning (p<0.05 in SVC at [-27,-4,1])^32–34^. We also tested for the integrated value signal, expressed as a sum of the value estimates of the model-based and model-free systems weighted by the arbitration control signal (P_MB_). The ventromedial PFC was positively correlated with the difference between the integrated value signals for the chosen and unchosen actions (p<0.05 in cluster-level corrected), fully consistent with previous reports on choice values^35–37^ (Figure 4, Supplementary Table S4).

Unlike the previous arbitration hypothesis^25^, our computational model also predicted that arbitration control influences how integrated values are converted into actual choices. Finally, we attempted to identify the brain regions involved in value–action conversion. We found that the inferior parietal lobe, insula (p<0.05 in FWE corrected), middle frontal gyrus, globus pallidus, FPC, supplementary motor area, and thalamus (p<0.05 in cluster-level corrected) are positively correlated with the probability value of the chosen action, referred to as the output value of the softmax function^38^. This finding is consistent with previous findings indicating stochastic action selection^39^. Other brain areas, such as the orbitofrontal cortex, superior temporal gyrus, middle frontal gyrus, supramarginal gyrus (p<0.05 in FWE corrected), medial PFC, superior frontal gyrus (p<0.05 in cluster-level corrected), posterior medial cortex, and lateral PFC, are negatively correlated with the chosen action probability. The negative encoding of stochastic action selection in the posterior medial cortex and the lateral PFC also replicates previous findings, as these regions have been implicated in the valuation of counterfactual choices^35,39,40^ (Figure 4, Supplementary Table S5).

### Effects of Depression on Goal-directed and Habitual Learning

To fully explore how depression affects the neural computations underlying model-based and model-free RL, we examined the relationship between the individual depression score and neural representations in each brain region implicated in model-based and model-free RL. We found evidence indicating the effect of depression on RL in multiple brain areas encoding prediction errors. The correlation coefficient between left insula activation and the SPE ([-36,20,-4], z=4.49), which represents the efficiency of neural encodings of the SPE in the left insula, was inversely proportional to the depression score (estimated correlation coefficient=-0.396, *p*=0.037; Figure 5a). We also found a significant negative correlation between the neural efficiency for encoding the RPE in the bilateral caudate (left: [-4,6,-4], z=4.50, right: [4,8,-4], z=5.30) and the individual depression score (estimated correlation coefficient=-0.412/-0.376, *p*=0.029/0.049 for the left and right, respectively; Figure 5b).

**Figure 5:**
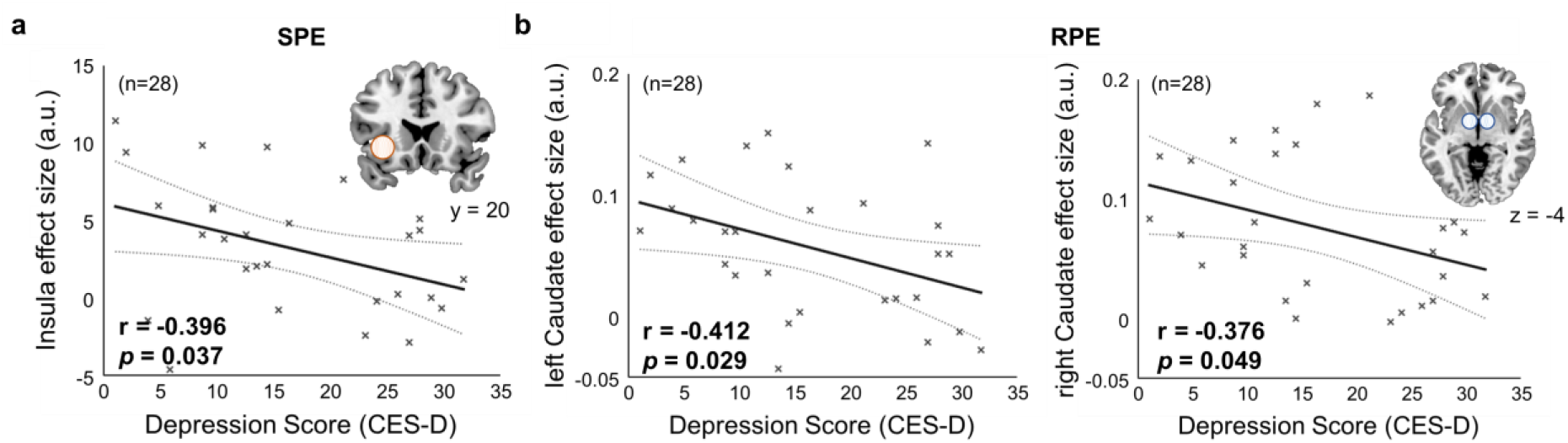
Parametric effect of depression on goal-directed and habitual learning. (a) Depression impacts on neural encoding of SPE information (n=28). The shaded circles represent seed regions for which parameter estimates of the GLM analysis were extracted. The seed region is the left insula, and the parameter estimates were extracted from our GLM analysis which regressed the SPE signal against the BOLD response ([-36,20,-4], z-score=4.49; Figure 4). The estimated effect size for the left insula is negatively correlated with the depression score. (b) Depression impacts on RPE response (n=28). The estimated effect size of RPE for bilateral caudate (left: [-4,6,-4], z-score=4.50, right: [4,8,-4], z-score=5.30) is negatively correlated with the depression score. a.u. stands for arbitrary units.

These findings directly demonstrate how depression affects value updates for goal-directed and habitual learning.

### Effects of Depression on Prefrontal Arbitration Control

We also tested for the neural effects of depression on arbitration control. First, we found that the individual depression score is significantly correlated with the learning rate for MF reliability estimation, the key variable required to quantify the reliability of predictions made by the model-free RL strategy based on the RPE (estimated correlation coefficient=0.335, *p*=0.007; Figure 6a, left; Supplementary Figure S1). This offers a theoretical prediction that depression entails over-sensitivity to the RPE, making arbitration control more sensitive to the prediction of the model-free system (Figure 6a, right). This implies that the brain regions implicated in mediating arbitration control focus on information about the reliability of the model-free system, rather than encoding the reliability information of the RL system controlling behavior at the moment (i.e. max reliability), leading to the disruption of neural computation underlying normal arbitration control.

**Figure 6:**
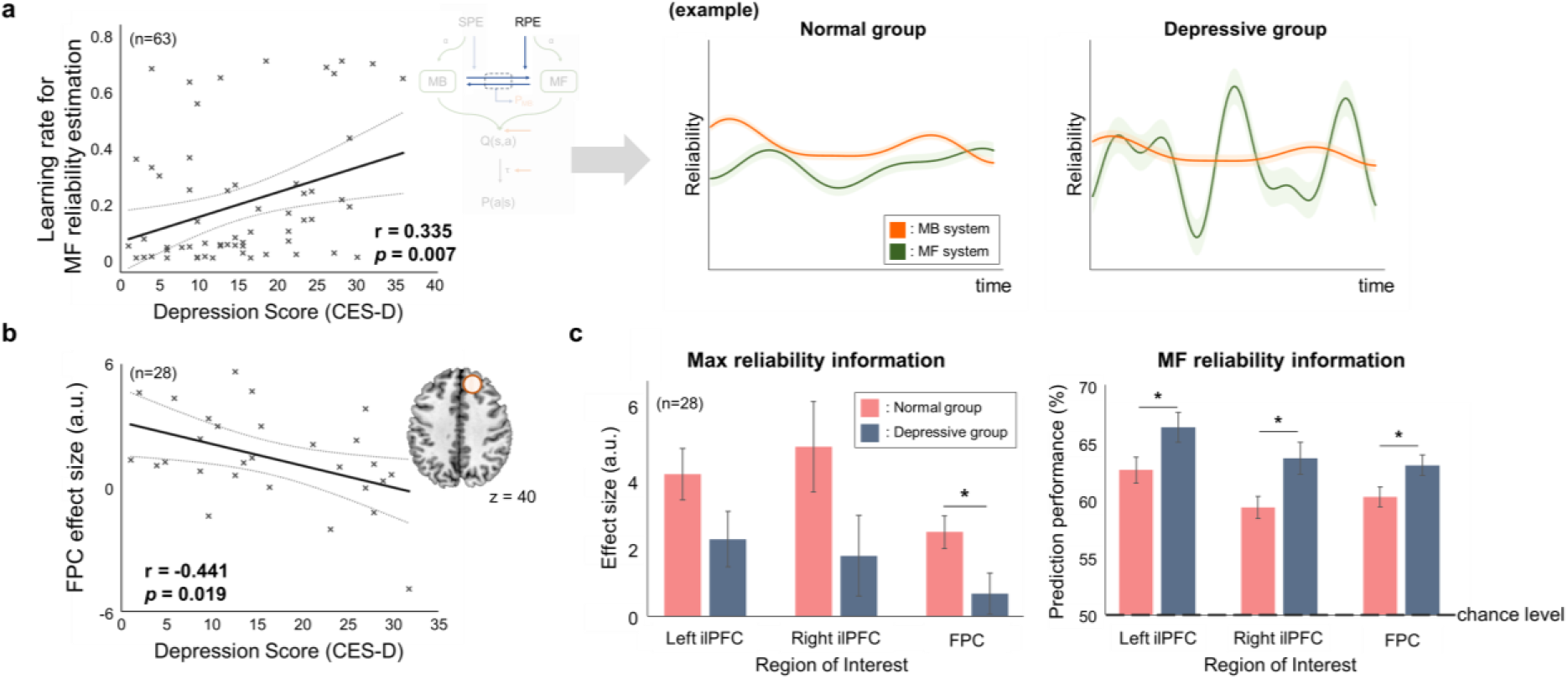
Parametric effect of depression on prefrontal arbitration control. (a) Depression effects on the model-free system’s sensitivity to RPE (n=63; the total number of subjects, including 40 who participated behavioral experiment only and 23 who were also scanned with the fMRI). (Left) Relationship between the depression score and the learning rate parameter for reliability estimation of the model-free system. Individuals with a higher depression score tend to exhibit a higher learning rate, indicating that their reliability estimation for the model-free system is more sensitive to RPE. (Right) Illustrative examples of reliability changes of people with a low (“normal group”) and a high depression score (“depressive group”), each of which is associated with a low and a high learning rate for reliability estimation, respectively. The depressive group shows rapid changes in MF reliability due to higher learning rate. (b) Relationship between the depression score and the estimated effect size of Max reliability for frontopolar cortex (FPC), the brain area implicated in arbitration control (n=28; the total number of subjects scanned with fMRI). The estimated effect size of the Max reliability signal for FPC (coordinates [8,44,40]; z-score=3.83) is negatively correlated with the depression score. (c) Comparison of reliability signal representation performance in normal (n=15) and depressive (n=13) group (GLM and MVPA analysis). The parameter estimates of the Max reliability for the bilateral ilPFC ([-52,26,16], z-score=4.48; [42,20,-8], z-score=3.61) and FPC ([8,44,40], z-score=3.83) tend to decrease in the depressive group. The MVPA analysis with these three seed regions reveals that the amount of information about the model-free reliability was significantly higher in the depressive group in all three regions (one-way ANOVA). Asterisk (*) indicates significant difference at the 0.05 level.

To test the prediction that depression disrupts neural processing pertaining to arbitration control, we conducted a GLM analysis with the max reliability signal. We found that the effect size (parameter estimates from the GLM analysis) of the max reliability signal for FPC ([8,44,40], z=3.83) was negatively correlated with the depression score (estimated correlation coefficient=-0.441, *p*=0.019; Figure 6b). Moreover, the effect sizes of the max reliability signal for the bilateral ilPFC ([-52,26,16], z-score=4.48; [42,20,-8], z-score=3.61) and FPC ([8,44,40], z-score=3.83), the brain areas previously implicated in arbitration control^25,29,41^, tended to be lower in the depressive group (CES-D score≥16) than in the control group (CES-D score<16) (Figure 6c, left; one-way ANOVA; F_1,26_=2.76 [*p*=0.108], F_1,26_=2.93 [*p*=0.099], F_1,26_=5.18 [*p*=0.031] for the left and right ilPFC and FPC, respectively).

To further evaluate the prediction that depression makes neural processing for arbitration control more sensitive to the model-free system, we ran an MVPA for the bilateral ilPFC and FPC. This analysis quantifies the amount of information concerning the reliability of the model-free system embedded in these brain areas. We used a support vector machine, an optimal neural network for prediction and generalization, to conduct a binary classification of model-free reliability (high vs. low; upper/lower 33^rd^ percentile threshold) and compared the prediction performance of the control and depression groups.

We found that the prediction performance of model-free reliability in the bilateral ilPFC and FPC was significantly higher in the depression group than in the control group (Figure 6c, right; one-way ANOVA; F_1,26_=4.57 [*p*=0.042], F_1,26_=4.33 [*p*=0.047], F_1,26_=4.89 [*p*=0.036] for the left and right ilPFC and FPC, respectively). On the other hand, the same analysis found no significant inter-group differences in model-based reliability signal or max reliability signal (Supplementary Table S6). Taken together, these results strongly support our arbitration control hypothesis that depression is associated with increased sensitivity to RPE, leading to instable arbitration in which the reliability of predictions of the model-free system becomes predominant and the reliability of predictions of the model-based system becomes less influential.

### Effects of Depression on Value-Action Conversion

Our computational model also explains how the degrees of control allocated to the model-based and model-free systems influence how value is converted into actual choice (exploitation sensitivity). For example, exploitative and explorative choices are associated with high and low exploitation sensitivity, respectively. We found that the exploitation parameter for model-based RL is negatively correlated with the individual depression score (correlation coefficient=-0.412, *p*=0.001; Figure 7a, left), indicating that subjects with higher depression scores exhibit more exploratory choices when their choices are guided by model-based RL (Figure 7a, right).

**Figure 7:**
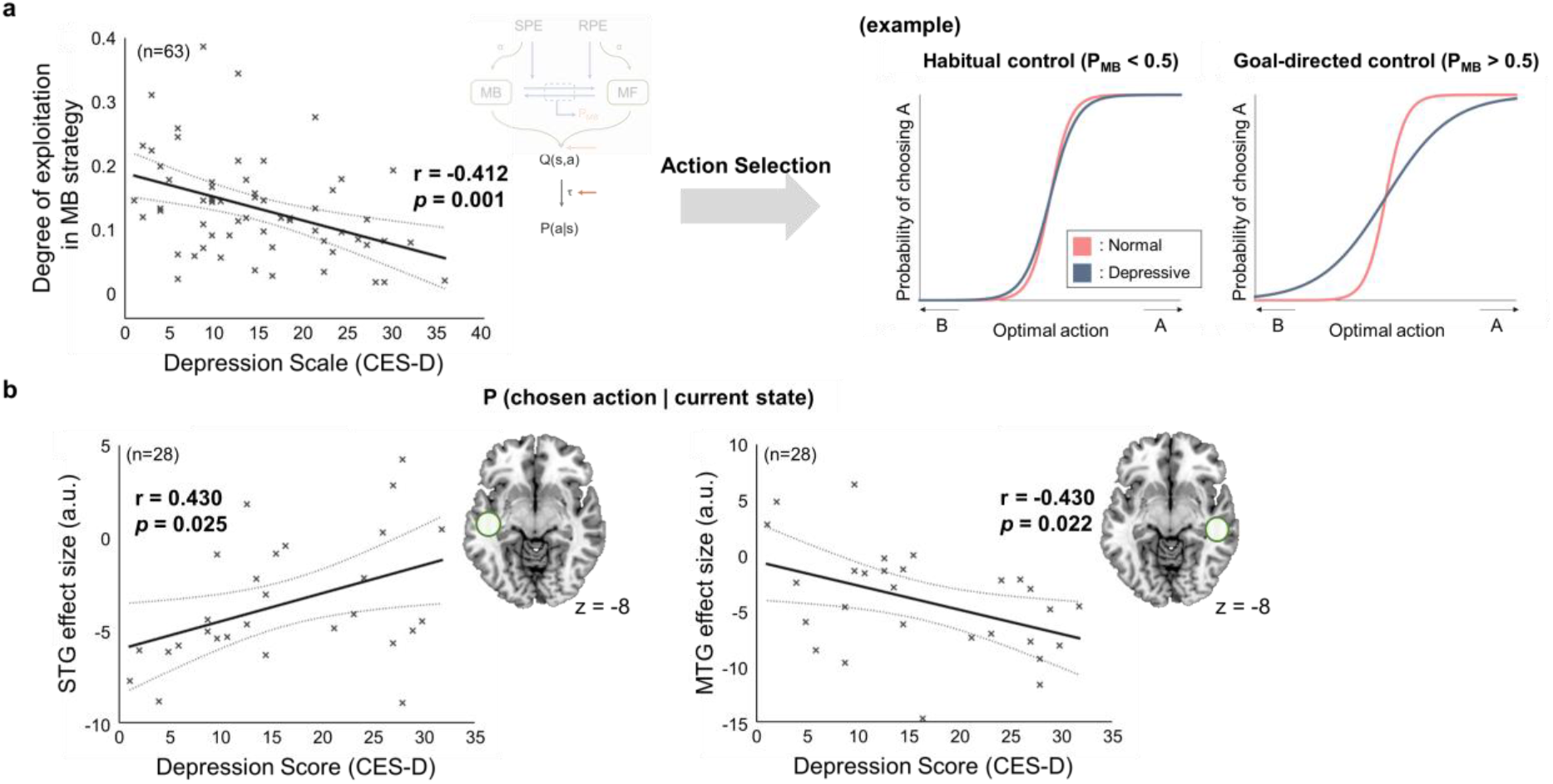
Parametric effect of depression on value-action conversion. (a) Depression effects on model parameters (n=63; the total number of subjects, including 40 who participated behavioral experiment only and 23 who were also scanned with the fMRI). (Left) Relationship between the depression score and the degree of exploitation in MB strategy (τ_MB_). The MB exploitation parameter decreases as the depression score increases. (Right) Examples illustrating exploitation parameter effects on action selection. Shown are the softmax functions that convert an action value into a choice probability value, between the normal and the depressive group for goal-directed and habitual control. Compared to the normal group (pink), the depressive group makes more exploratory choices especially as they rely more on the model-based system (blue). (b) Relationship between the depression score and the parameter estimate of the probability of selecting chosen action for the two seed regions, right Superior Temporal Gyrus (STG) ([-46,-20,-8]; z-score=4.21) and the left Middle Temporal Gyrus (MTG) ([56,-32,-8]; z-score=3.55). The effect size of right STG and left MTG increases and decreases with the individual severity of depression, respectively.

In the subsequent neural analysis, we explored the relationships between the depression score and the neural representations of value–action conversion. The parameter estimates of the probability value of taking the chosen action are significantly correlated with the depression score in two brain regions. The parameter estimates in the two seed regions—the right superior temporal gyrus (STG; [-46,-20,-8], z=4.21) and left middle temporal gyrus (MTG; [56, 32, -8], z=3.55)—are positively and negatively correlated with the CES-D score, respectively (correlation coefficient=0.430, *p*=0.025 for STG; correlation coefficient=-0.430, *p*=0.022 for MTG; Figure 7b).

## DISCUSSION

By combining a computational model allowing for sub-optimality in learning and decision-making, a model-based fMRI analysis, and an MVPA, the present study fully characterizes how depression influences the different levels and stages of RL: value computation, prefrontal arbitration control for value integration, and value–action conversion. We found that depression has a parametric effect on neural representations of prediction error for model-based and model-free systems, respectively, explaining how depression hampers value computation ability. Another intriguing finding is that the brain areas implicated in arbitration control, bilateral ilPFC and FPC, become more sensitive to the predictions of the model-free system in people with depression, indicating that depression disrupts the balance between goal-directed and habitual control. We also found that depression increases the tendency to make exploratory choices during model-based control, but not during model-free control.

### Computational Theory of Prefrontal Goal-directed and Habitual Control

The present study’s computational model of dynamic arbitration control of model-based and model-free RL allows us to explore the full parametric effects of depression on prefrontal goal-directed and habitual control. Although mounting evidence suggests that prediction uncertainty might be a key variable for prefrontal arbitration control^6,25,27^, little is known about the computational reasons people with depression tend to exhibit behavioral biases towards either goal-directed or habitual behavior. Addressing this issue involves a few challenges. First, simply evaluating the two separate hypotheses contradicts the prevailing view that the brain circuitries guiding goal-directed and habitual behavior interact with each other. Second, there is no guarantee that a rational arbitration control model is flexible enough to explain the individual variability associated with depression. Third, exploring a depression-specific model based on the assumption that depression follows a computational regime that substantially deviates from rational decision-making may enable us to explain severe depression, but cannot explain a continuum extending from a normal to a severely depressed state.

To fully address these issues, we considered a computational model of dynamic arbitration control allowing for individual variability in suboptimal learning and decision making. Intriguingly, we found that the influence of prediction uncertainty is not confined to value integration^25^, but extends as far as value–action conversion. This also allowed us to test the full effect of depression on decision-making at different computation levels: model-based and model-free reinforcement learning, arbitration control for value integration and value–action conversion.

### Effects of Depression on Neural Representations of Prediction Error

Our study found that depression has a parametric effect on the neural representations of the two distinct types of prediction errors associated with model-based and model-free RL: SPE and RPE. The neural analysis revealed that depression scores were correlated with an attenuation of the SPE signal in the left insula and the RPE in the bilateral caudate.

Dopamine is crucial for both model-free and model-based RL. Numerous previous studies have reported dopamine’s role in guiding the RPE^42,43^, and a recent finding discussed the essential role of dopamine in stimulus-stimulus associative learning^44^, implicating the involvement of dopamine in SPE representation. Depression is characterized by decreased dopamine levels^45,46^, which may impair learning in both model-free and model-based systems. In fact, RPE signals in depression have reportedly been reduced in various experimental conditions (both Pavlovian learning^12^ and instrumental learning^13–17^). Our study not only corroborates previous findings concerning RPE deficits in depression, but also further suggests that depression may impact neural representations of SPE.

### Effects of Depression on Prefrontal Arbitration Control

One interesting prediction of the model is that CES-D is positively correlated with the parameter value for controlling the learning rate for updating model-free reliability based on the RPE, indicating that reliability estimation for the model-free strategy is very sensitive to RPE changes. This suggests that, in people with high CES-D scores, arbitration control might be predominantly driven by the model-free reliability signal, rather than by a fair comparison of the model-free and model-based reliability signals. We explored this possibility through a combination of a general linear model (GLM) and MVPA.

Our GLM analysis showed the negative effect of the depression on the neural representations of arbitration control in the prefrontal cortex. In bilateral inferior lateral PFC and frontopolar cortex, the brain areas reportedly encoding the key variable for arbitration control^25^, neural representations tend to be weaker in the high CES-D score group. Critically, the subsequent MVPA shows that the amount of reliability information of the model-free system is significantly higher in the high CES-D score group. Taken together, these findings theoretically implicate that depression may hinder the PFC’s ability to estimate the reliability of each learning strategy from the corresponding prediction error.

### Effects of Depression on Valuation-Action Conversion

The present study also provides a computational and neural account of how depression causes sub-optimal action selection. Our computational model predicts that depression increases the tendency to make exploratory choices during model-based control, rather than model-free control.

This finding could also clarify the two conflicting views of choice consistency behaviors in depression^47,48^. Beever et al. (2013) found no significant difference in exploration pattern in a reward-maximizing task between a normal and a depressed group. Blanco et al. (2013), on the other hand, found that a depression group tended to explore more. This conflict might be attributable to differences in task structure. Beever et al.’s study used a task with a relatively stable environmental structure, such that people performed tasks relying on the model-free system. This is consistent with our view that depression has a relatively weak influence on exploration during model-free control. However, the reward structure used in Blanco’s study encouraged more frequent policy changes, accommodating the need for model-based control. This is also consistent with our view that exploratory choice behavior becomes more pronounced during model-based control.

The neural results of the present study, which show that the STG response is higher in people with depression, are fully consistent with previous finding that STG response increases when people switch to other options rather staying^49^. Our results address not only the implication for the role of STG in exploration, but also how depression influences exploration at the neural level.

Our finding that the degree of exploitation decreases as CES-D score increases (shown in Figure 7) explains why reward sensitivity is reduced in people with depression. A decreasing degree of exploitation decreases the tendency to convert a learned policy into an actual choice, reducing the efficiency of translating changes in the reward structure into changes in actual choices. This is also consistent with the view that our model’s degree of exploitation parameter can be interpreted as reward sensitivity^16,50^. In addition to supporting existing evidence of declined reward sensitivity in depression^51–53^, the present study advances the view by proposing that this tendency becomes stronger during model-based control.

### Potential clinical applications

The present findings suggest how depression influences goal-directed and habitual control in the prefrontal–striatal circuitry. The full characterization of the effects of depression on different stages of learning and decision-making creates possibilities for various clinical applications. First, our neural results explain why such brain stimulus techniques as repetitive transcranial magnetic stimulation (rTMS) and deep brain stimulus (DBS) to the frontopolor cortex^54^ or ventral striatum^55^ are effective in alleviating depressive symptoms. Second, our theoretical idea suggests that behavioral therapy to reduce sensitivity to reward prediction errors might help people with depression regain a balance between goal-directed and habitual control. Intriguingly, our findings also indicate a parametric effect of depression on learning and decision-making in a relatively young age group (average=22.8 yrs). Considering the onset of MDD is approximately 25 to 45 yrs^56^, our study offers possibilities for not only investigating how mild depression transitions to MDD, but also developing clinical applications for the early diagnosis of MDD.

## METHODS

### Participants

Sixty-five right-handed Koreans (28 females; mean age of 22.8±3.8) participated in the study. Participants were recruited from the local society through the online announcement. Only 28 subjects were scanned with fMRI during the task. Two subjects whose total accumulated reward are below the chance-level (mean amount of rewards with 10,000 random simulations) were excluded from the analysis. Thus, a total of sixty-three behavioral data and twenty-eight fMRI data were left for the analysis. To acquire the depressive level of individuals, people were instructed to complete the Center for Epidemiologic Studies Depression (CES-D) questionnaires^26^ before the experiment (For the distribution of participant’s depression severity, see Supplementary Figure S2).

No subjects had the history of neurological of psychiatric disease. Every subject provided written consent to the experimental protocols which were approved by the Institutional Review Board (IRB) of the Korea Advanced Institute of Science and Technology (KAIST).

### Task

We used the sequential two-stage Markov decision task proposed to dissociate model-based and model-free learning strategy^25^. In this task, subjects make a binary choice (either left or right) and proceed to the next state with a certain probability. When the next state is appeared in the screen, participants make another choice. The two consecutive choices is followed by a transition to an outcome state. Subjects perform 100 trials in the pre-learning session to learn the structure of the task. Four main sessions with 80 trials on average follow the pre-learning session. Participants are instructed to collect as many coins as possible in the main sessions.

The task consists of two conditions: a specific-goal condition and flexible-goal condition. The goal condition is indicated by a color of a box at the beginning of each trial. In the specific-goal condition, participants are presented with a box with a specific color (red, blue, yellow). A monetary reward is given only when the coin color matches with the color of the given box. In the flexible-goal condition, on the other hand, participants are given a white coin box with which all types of coins become redeemable. Two types of state-transition probability are used to control the uncertainty of the environment. The state-transition probability (0.5, 0.5) and (0.9, 0.1) is intended to implement the highly-uncertain and relatively less uncertain environment, respectively. The four types of block (2 goal-conditions x 2 uncertainty conditions) are presented in pseudo-random order. Each block consists of 4-6 trials.

### Computational Models

The computational model of this study is motivated by the previous arbitration control hypothesis that prediction uncertainty of the model-based and model-free RL is a key variable to guide value integration of the two corresponding systems^25^. The model consists of the three processes: value learning, arbitration, and action selection (Figure 3a).

In the value learning stage, both a model-based and model-free system learn action values for each state. A model-based system uses state prediction error (SPE = 1-expected transition probability) to update the state-action-state transition probability, by using a FORWARD learning^27^ and learns action values by combining the learned state-action-state transition probability and reward in the outcome state. For a model-free system, on the other hand, the state-action value learning is based on RPE (RPE = actual value-expected value). It is implemented with a SARSA algorithm^5^.

In the arbitration process, the reliability estimation of the model-based system was implemented with a hierarchical empirical Bayes method using the history of the SPE, the reliability of the model-free system was implemented with the Pearce-hall associability rule using an unsigned RPE. These estimated reliability signal were then used to guide the competition between the two systems, which is implemented with a dynamic two-state transition model. The output of this model is a model choice probability (P_MB_), used as the control weight for value integration of the two systems. Finally, in the action selection stage, the model selects the action stochastically using softmax rule^38^. For more details, refer Lee et al (2014).

In this study, we suggested two variants of arbitration control, allowing for sub-optimality in value learning, arbitration, and action selection: one version with separate model-based and model-free learning and another version with a dynamic exploitation. The former type of the model assumes the different learning rates of a model-based and a model-free system. The latter class of models is based on the former model, with the further assumption that the degree of exploitation, an indicator of optimality of the RL agent’s policy, is a function of the model choice probability, P_MB_. We tested three different types of exploitation as follows: logistics, linear, weighted linear. Note that in all cases, we set the parameters of the model in a way that is reduced to the original RL with a single exploitation parameter.

### fMRI Data Acquisition

Functional imaging was conducted on a 3T Siemens (Magnetom) Verio scanner located in the KAIST brain imaging center (Daejeon). Forty-two axial slices were acquired with interleaved-ascending order at the resolution of 3 mm x 3 mm x 3 mm, covering the whole brain. A one-shot echo-planner imaging pulse sequence was utilized (TR = 2800 ms; TE = 30 ms; FOV = 192 mm; flip angle = 90°). The high resolution structural image was also acquired for each subject to the resolution of 0.7 mm X 0.7 mm x 0.7 mm.

### fMRI Data Pre-processing

Images were processed and analyzed using the SPM12 software (Wellcome Department of Imaging Neuroscience, London, UK). The first two volumes were removed to reduce T1 equilibrium effects. The EPI images were corrected for slice timing, motion movement and spatially normalized to the standard template imaging provided by SPM software.

For the general linear model analysis (GLM), normalized images were smoothed with 6mm FWHM Gaussian Kernel and a high-pass filter (128s cut-off) was applied to remove the noise.

For the multivoxel pattern analysis (MVPA), unsmoothed EPI image data was used. Detrending and z-scoring were processed to reduce the linear trends and to match the range of the signal.

### General Linear Model Analysis (GLM)

Subject-specific value-related signals and arbitration control signals were computed from the arbitration model, and the signals were regressed against voxel-wise signals from the EPI image set. The order of the regressors is as follows: prediction error from the model-based system (SPE) and the model-free system (RPE), reliability comparison signal which is a key variable for arbitration control (=max (Rel_MB_, Rel_MF_); max reliability), the chosen value of model-based system (Q_fwd_), the chosen value of model-free system (Q_sarsa_), the difference between chosen and unchosen integrated values (Q_arb_) and the probability of selecting chosen action (P_chosen action_). The regressors were serially non-orthogonalized in the GLM analysis to prevent the effect of regressor orders in the interpretation of the results. MARSBAR software (http://marsbar.sourceforge.net) was used to extract parameter estimates from the region of interest^57^.

### Multivoxel Pattern Analysis (MVPA)

The MVPA analysis was conducted to quantify types and amounts of information encoded in specific the region of interest (ROI). The classification performance is regarded as the amount of information pertaining to the variable of interest. Three ROIs, left/right inferior lateral prefrontal cortex (ilPFC) and Frontopolar prefrontal cortex (FPC), were selected, which were known to engage in the arbitration control process. Masks of each brain region were functionally defined from the GLM analysis. We used the clusters whose response to the Max reliability signal survived after the whole-brain correction (*p*<0.001, uncorrelated) as a mask for each ROI. We set the BOLD response time 4-6s.

A binary Support Vector Machine (SVM) classifier was applied to learn voxels patterns with each ROI. For each subjects data, the SVM was trained to best match its output to a binarized reliability-related signal (MB reliability, MF reliability, or Max reliability); the 33^th^ and 67^th^ percentile threshold were used to define the two classes, ‘high value group’ and ‘low value group’, respectively. All voxels in the mask were used for learning. The input dimension is 196, 79, 294 for left ilPFC, right ilPFC and FPC, respectively. The average number of data from each subject are 350 for MB reliability, 549 for MF reliability and 543 for Max Reliability. Thirty-fold cross validation was conducted for evaluation. All processes were implemented based on the Princeton Multi-Voxel Pattern Analysis toolbox^58^. Finally, an ANOVA analysis was conducted to compare signal prediction accuracy between the normal and depression group; the two subject groups were defined by using the standard cutoff criteria of CES-D score, 16^59,60^.

### Data Availability

All data analyzed in this study will be available upon request.

## ACKNOWLEDGEMENTS

This research was supported by the Brain Research Program through the National Research Foundation of Korea (NRF) funded by the Ministry of Science, ICT & Future Planning (NRF-2016M3C7A1914448), NRF funded by the Korea government (MSIT) (No. NRF-2017R1C 1B 2008972), the research fund of the KAIST (Korea Advanced Institute of Science and Technology) (Grant code: G04150045), and Institute for Information & Communications Technology Promotion (IITP) grant funded by the Korea government (No. 2017-0-00451).

## Author Contributions

S.H. and S.W.L. designed the study, analyzed the data and wrote the manuscript. S.H. conducted the experiments.

